# Receptor-substrate competition for the TonB homologue FusB suggests a model for ferredoxin import in *Pectobacterium* spp

**DOI:** 10.1101/2025.10.16.682781

**Authors:** Marta Wojnowska, Victor Flores, Tamas Yelland, Stuart R. Fisher, Agnieszka Bogucka, Katherine Stott, Daniel Walker

## Abstract

TonB-dependent uptake systems of Gram-negative bacterial pathogens constitute prominent virulence factors, allowing nutrient acquisition, primarily siderophore-bound iron, to cross the highly impermeable outer membrane (OM). Remarkably, the ferredoxin uptake system (Fus) of *Pectobacteriaceae*, a group of soft rot-inducing plant pathogens, imports an entire folded host protein into the periplasm and extracts its bound iron for growth. The inner membrane protein FusB, a TonB homologue, plays two roles in facilitating ferredoxin import. First, like other TonBs, it remodels the globular plug domain obstructing the lumen of the OM receptor FusA to allow ferredoxin passage. Unusually for a TonB protein, FusB then interacts directly with the FusA-bound ferredoxin substrate to facilitate its transport into the periplasm. In this work, we describe structures of FusB-ferredoxin and homodimeric FusB complexes and determine the key features of the binding interfaces formed by FusB with FusA and ferredoxin. We postulate that under resting conditions FusB exists a homodimer, stabilised by an intermolecular R241-D322 salt bridge. The homodimer dissociates when the “FusB-box” of FusA outcompetes one protomer, and FusA D53 displaces FusB D322. Upon ferredoxin binding, FusB undergoes a structural rearrangement, expanding its β-sheet from three to four strands. In agreement with the proposed sequence of events, ferredoxin binding displaces the receptor (FusA) from FusB with Arg241 forming an intramolecular salt bridge with Asp322 to stabilise the newly formed β-hairpin of FusB. We propose a mechanistic model for ferredoxin import in which FusB Arg241 acts as a molecular switch, and two distinct regions function as interaction hotspots.

## INTRODUCTION

Nutrient piracy is one of the major forces driving host-pathogen coevolution. Bacterial pathogens have developed diverse strategies to hijack nutrients necessary for growth from their host, particularly iron which is essential for the function of numerous enzymes^1,2^. Competition for iron has been shown to underpin microbial interactions in the rhizosphere community, with a dramatic impact on bacterial plant pathogens^3^. In Gram-negative bacteria the outer membrane is a formidable barrier against most bactericidal agents, however it also necessitates active import of scarce but essential nutrients into the cell^4–6^. The first step - import across the outer membrane into the periplasm – is supported by TonB-dependent uptake systems, which constitute prominent virulence factors. These pathways are mostly iron-specific: the metal ion is first tightly bound by a secreted siderophore, an iron chelator, and then delivered to a dedicated outer membrane receptor for import into the cell. Blocking or hijacking TonB-dependent uptake systems to deliver antibiotics is a promising antimicrobial strategy, with Trojan horse antibiotics such as sideromycins being of particular interest^7,8^.

Nutrient uptake into the periplasm requires a dedicated TonB-dependent receptor (TBDR) in the outer membrane and a complex of three proteins in the inner membrane: TonB, ExbB and ExbD^4–6,9^. The latter two form a motor complex that harnesses energy from proton motive force (PMF), while TonB, anchored in the inner membrane, stretches across the periplasm with a long proline-rich linker leading into a globular C-terminal domain (CTD). The receptor is a large 22-stranded β-barrel with the lumen occluded by an N-terminal globular plug domain. Nutrient binding on the extracellular face of the receptor triggers conformational changes that result in the release of a short unstructured portion of the plug domain into the periplasm, where it can interact with TonB-CTD. Within this unstructured region is the so-called TonB-box, with high β-strand propensity, which aligns against the third β-strand of TonB-CTD. Although the reported affinities of TonB for TonB-boxes of various receptors are often in the micromolar range^10–14^, the interaction via strand addition has been shown to be mechanically strong, allowing TonB to transduce the PMF-generated force to extract the force-labile portion of the receptor plug^14^. The mechanistic details of TonB-mediated plug remodelling remain uncertain, although it is thought that the pulling force is exerted by a rotary movement of TonB, coupled to the rotation of the ExbBD motor^6,15,16^.

The number of TBDRs in Gram-negative bacterial pathogens varies, with some species predicted to encode over 100 different TBDRs^17^ . In *E. coli* this number varies from 9 (strain K12) to 18 (strain O157:H7), and all of these are serviced by one TonB^4,6^. Some bacteria, however, have multiple TonB proteins encoded in their genomes, which tend to be receptor-specific^181911^. In the context of iron piracy some siderophore-independent systems have evolved, including the ferredoxin uptake system (Fus) found in soft rot plant pathogens of *Pectobacteriaceae* family. Fus is unusual because it imports a folded protein, plant host ferredoxin, into the periplasm, where a dedicated protease, FusC, cleaves it to release its 2Fe-2S iron-sulphur cluster^11,20,21^. Plant ferredoxins are small (∼10 kDa) proteins involved in a range of metabolic pathways and their different isoforms are found in the tissues of higher plants^22^. FusB appears to have two roles: remodelling the plug domain of the FusA receptor, and then directly engaging the ferredoxin substrate to facilitate its translocation into the periplasm^11^. This most likely entails two separate rounds of force transduction, energised by the PMF, and we have previously demonstrated that substrate interaction requires at least one of the two consecutive arginines (R241 and R242) of FusB that replace the canonical QPQ motif of TonB. Notably, among 5 TonB proteins (excluding FusB) present in *Pectobacterium carotovorum* LMG2410 (*Pc*LMG2410) the one most closely related to *E. coli* TonB is essential for generic iron import via canonical TBDRs, but not ferredoxin uptake^11^.

In this study we probed ferredoxin uptake mechanism further - using structural and biophysical approaches we characterised three distinct binding modes of FusB-CTD, with a dynamic interaction hotspot located on the third β-strand. This region interacts with the FusB carboxyl terminus either in *trans* (FusB homodimer) or in *cis* (ferredoxin-binding interface), however it also binds to the FusB-box of FusA. One key residue of FusB, R241, forms inter- or intramolecular ionic interactions with D322, however, when the FusB-box of FusA emerges in the periplasm, R241 switches to FusA D53 to facilitate plug remodelling. We have identified several FusB, FusA and ferredoxin residues contributing to the binding interfaces, and verified FusB R242 as critical for substrate interaction. These findings, along with assays on a range of protein variants, allowed the elucidation of a molecular switch mechanism involving FusB R241.

## RESULTS

### FusB-CTD dimerisation involves a salt bridge

To explore the key interactions involved in homodimer formation of FusB we obtained diffracting crystals of FusB-CTD and solved the structure to 2.08 Å by molecular replacement using monomeric *E. coli* TonB (*Ec*TonB) (PDB ID: 2GSK^23^). FusB-CTD forms a homodimer with a non-crystallographic 2-fold symmetry (Fig. 1A). The DALI server^24^ identified the closest fold in terms of structure to be that of *E. coli* TonB (PDB ID: 2GSK and 1U07). Similar to the structure of the 92-residue C-terminal construct of *E. coli* TonB^25^ (PDB ID: 1U07), FusB-CTD monomers form an intermolecular antiparallel β-sheet, resulting in a buried surface area of 2046 Å^2^ as determined by the PISA server^26^. An intermolecular salt bridge between R241 and D322 appears to staple each C-terminal region to the adjacent monomer. R241 is one of two consecutive arginines that replace the canonical QPQ motif found in TonB proteins and we have previously reported that substitution of the double arginine motif (R241, R242) in FusB with lysines abolishes import of the ferredoxin substrate^11^.

**Figure 1.**
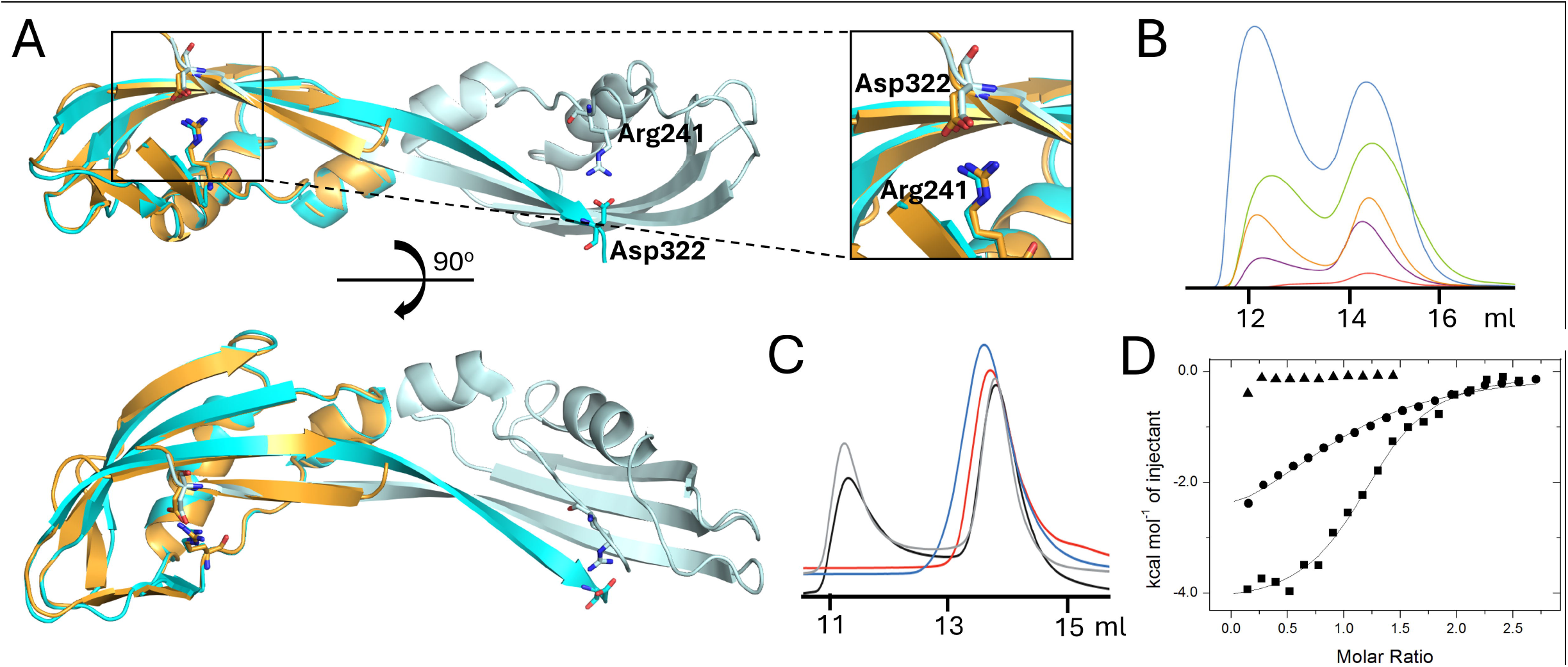
FusB-CTD dimerization and key role of R241-D322 salt bridge. A – Crystal structure of the FusB-CTD dimer (cyan and pale cyan) and AlphaFold3-predicted monomer (gold) overlaid onto the cyan molecule. R241 and D322 forming the intra- or intermolecular salt bridge are shown as sticks. B – size exclusion chromatograms of FusB-CTD at varying concentrations: 0.2 mg loaded - red, 0.5 - purple, 0.8 - orange, 1.2 - green and 2 - blue. C – size exclusion chromatograms of *Pc*FusB- CTD wild type (black) R241K (red), R242K (grey) and D322N (blue); 0.5 mg protein was loaded. D – Isothermal calorimetry titrations of SpFer into FusB-CTD R241K (circles), R242 (triangles) and D322N (squares).

Interestingly, the prediction of a FusB-CTD monomer by AlphaFold3 indicated that the extended C-terminal strand folds over, resulting in a 4-stranded b-sheet (Fig. 1A, gold), where the terminal strand is anchored via the same the same R241-D322 salt bridge, in this case intramolecular. We have in fact observed that FusB-CTD elutes as two distinct species in size-exclusion chromatography and the proportion of the dimer species increases with protein concentration (Fig. 1B). It therefore seems likely that the monomer-dimer equilibrium of FusB involves *cis* and *trans* switching of the R241-D322 salt bridge.

### Double arginine motif: R241 is critical for dimerization, R242 for substrate binding

The structure of FusB-CTD indicates that R241 is involved in the formation of the salt bridge with D322, which appears to stabilise the FusB dimer. Since R242 does not seem to play a role in dimerization we hypothesised that it may play a key role in substrate binding. We therefore generated relevant site-specific variants to test these hypotheses. The single R241K and D322N variants eluted from gel filtration as a single monomeric species (Fig. 1C), whereas the R242K variant eluted as two species, similarly to wild-type FusB-CTD. The monomeric constructs (R241K and D322N) did not exhibit any signs of dimerisation even at high protein concentrations (Fig. S1). Both monomeric constructs interact with FusB-CTD, but while the affinity of D322N FusB-CTD for ferredoxin (K_D_ = 7.4 mM) is comparable to that of wild-type FusB-CTD (see Methods and Table S3), R241K FusB-CTD shows an approximate 3-fold reduction in affinity for ferredoxin (K_D_ = 25.5 mM). In contrast, titration of ferredoxin into R242K FusB-CTD produced almost no detectable signal (Fig. 1D, Table S3). These results imply that both arginine residues contribute to substrate binding, although to hugely varying extents. We have identified R242 as critical for ferredoxin interaction while R241, along with D322, is essential for FusB dimer formation. Furthermore, the involvement of the R241-D322 salt bridge in dimerization suggests that the interface observed in the FusB-CTD homodimer structure persists in solution and is not a crystallographic artefact.

### Structure of FusB-CTD in complex with ferredoxin

The apparent molar mass of the FusB-CTD-ferredoxin complex was determined by SEC-MALS as 24.3 kDa (Fig. S2) which is very close to the expected 24.6 kDa for a heterodimeric complex. As numerous attempts to co-crystallise FusB-CTD with various plant ferredoxins did not yield crystals of the complex, we employed AlphaFold3 to predict its structure. Although the resulting predictions almost always indicated the involvement of the critical residue R242 and often also R241 in complex formation, the modes of binding in the top scoring predictions differed substantially, involving different regions of the ferredoxin. Therefore, a co-crystallisation approach was undertaken, whereby FusB-CTD and ferredoxin were fused through a connecting flexible linker. The construct was designed such that it contained N-terminally His_6_-tagged FusB-CTD followed by a 12-residue flexible linker with SpFer at the C terminus; the tag was removed during purification.

This fusion construct yielded crystals in several different screens with the best crystals diffracting to 1.7 Å. The asymmetric unit contained one fusion protein, however the interface between FusB-CTD and ferredoxin within the asymmetric unit did not appear biologically relevant with contacts mostly mediated via water molecules, and the critical arginine motif of FusB is not involved. Conversely, the interfaces formed with the neighbouring symmetry-related molecules seemed plausible, with numerous hydrogen bonds, hydrophobic interactions and the engagement of the FusB double arginine motif (Fig. 2A and B). The FusB-ferredoxin binding interface is smaller (1423 Å^2^) than that in FusB-CTD dimer, which is probably due to the small size of the substrate and the fact that only a limited surface area of the ferredoxin translocating via the FusA lumen would likely be accessible to FusB. Importantly, the FusB-CTD monomer interacting with the substrate adopts the same conformation as in the AlphaFold3-generated monomer, whereby the long terminal b-strand folds over and the interaction between the resulting two shorter strands is stabilised by an intramolecular R241-D322 salt bridge. The DALI server^24^ identified *Ec*TonB as the closest fold for the monomeric FusB-CTD – this time a monomeric construct comprising 4 β-strands (PDB ID: 1XX3^12^).

**Figure 2.**
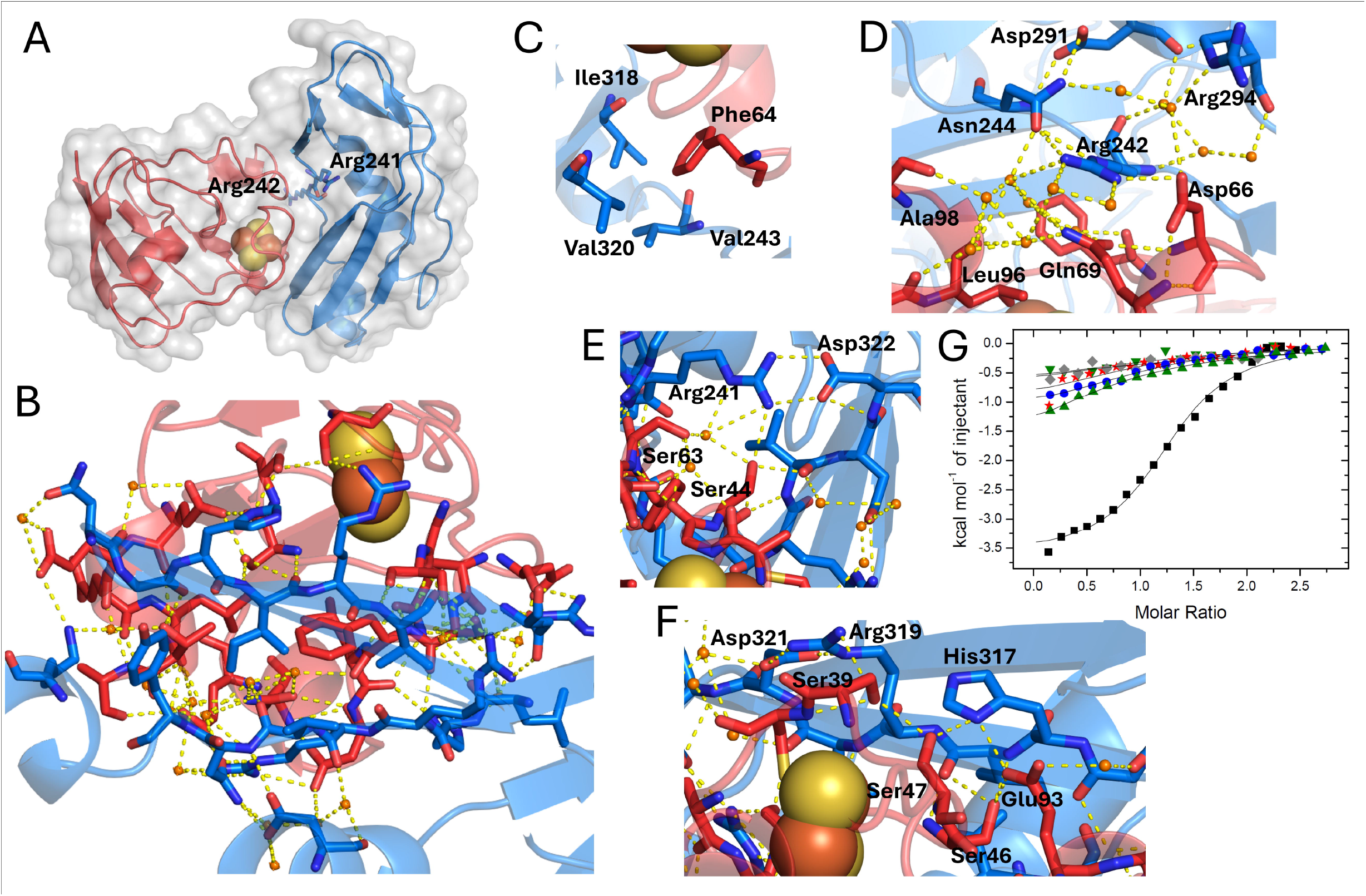
FusB-ferredoxin binding interface. A – FusB-ferredoxin complex with FusB-CTD shown in blue and SpFer in red; R241 and R242 are shown as sticks, iron-sulphur cluster as spheres. B – interfacing residues (shown as sticks) and water molecules (orange spheres) with polar contacts indicated with yellow dashed lines. C - F – close-up views of (C) SpFer F64 wedged into the hydrophobic pocket of FusB-CTD; D - FusB-CTD R242 interaction network; E - FusB-CTD R241-D322 salt bridge and SpFer residues interacting with R241; F - FusB-CTD H317, R319 and D321 interaction network. G and H: ITC thermograms from titrations of various ferredoxins and their variants into FusB-CTD. G - SpFer wild type (squares), F64Y (blue circles), D66N (grey diamonds), E93K (inverted green triangles), E93A/E94A (green triangles) titrations into FusB-CTD wild type as well as SpFer wild type titration into H317A (red stars).

The most striking feature of the co-crystal structure are several extensive networks of water molecules, which connect even more distantly positioned residues of both interacting proteins and are thus likely to contribute to stabilising the interaction (Fig. 2B). At the centre of the interface lies highly conserved SpFer F64, inserted into a hydrophobic pocket between the N-terminal unstructured region and the C-terminal strand of FusB (Fig. 2C). The critical FusB residue R242 is positioned on the edge of the interface; its side chain forms a hydrogen bond with SpFer D66 and potential pi-stacking interactions with Q69 (Fig. 2D). FusB N244 forms a water-mediated hydrogen bond with ferredoxin Q68 and, in addition, interacts with R242 side chain, on the opposing side to SpFer D66, which appears to fix R242 in place. R241 forms the aforementioned intramolecular salt bridge with D322 but its side chain also makes two contacts with the ferredoxin, one direct (S44) and one water-mediated hydrogen bond (S44 and S63) (Fig 2E). Two salt bridges are formed by FusB residues H317 (Fig. 2F) and L250, which interact with ferredoxin E93 and E94, respectively. In addition, Nδ1 of H317 interacts with the SpFer S47 side chain, while K250 is involved in coordination of a water molecule along with SpFer T97 hydroxyl and E94 backbone carbonyl. Another direct hydrogen bond is between the side chains of SpFer S39 and FusB R319; similar to R242, located on the other side of the interface, the R319 guanidine moiety is locked in place, in this case by a salt bridge with FusB D321 (Fig. 2F). A number of backbone interactions contribute to the binding interface, some of which involve the side chains of FusB R237 and H240 as well as SpFer Q62. Interestingly, most of the direct interactions involve positively charged residues on FusB side (3 arginines, 1 lysine, 2 histidines), while in ferredoxin it is the negatively charged residues (1 aspartate, 2 glutamates) that contribute to the binding interface, along with 6 serines located in the loops surrounding the iron-sulphur cluster.

### FusB-ferredoxin interface validation

The FusB-ferredoxin interface in the co-crystal structure is consistent with the effect of mutation of R241 and R242 on complex formation. Further support is provided by the fact that FusB D322, although important for salt bridge formation, is not involved in any interactions with the ferredoxin in the structure, and the D322N mutation did not affect binding to ferredoxin in ITC as described above. In addition, both monomeric and dimeric folds of FusB-CTD appear plausible based on the available TonB protein structures.

To further validate the FusB-ferredoxin interface we generated additional site-specific variants of each protein and tested these using ITC. Consistent with the co-crystal structure, the FusB-CTD H317A variant (H317 interacts with E93 and E94 of ferredoxin) exhibited severely reduced and indeterminable affinity for SpFer1 (Fig. 2G). A similar phenotype was observed with the SpFer1 variants D66N and E93K titrated into FusB-CTD, reflecting the importance of their respective interactions with R242 and H317 of FusB. Notably, the E93K mutation was previously reported to affect ferredoxin function by destabilising the loop which coordinates the iron-sulphur cluster^27^. As a charge switch is quite a dramatic change, and E94 of the ferredoxin also appears to be involved in FusB interaction, we generated a less disruptive, double alanine variant E93A/E94A, which exhibited approximately 4-fold reduction in affinity for FusB-CTD (K_D_ = 31.3 mM, Fig. 2G). In addition, we substituted SpFer1 F64 with a tyrosine to verify the importance of the central hydrophobic pocket; it appears that the introduction of a hydroxyl group has a considerable impact on binding enthalpy and affinity, reducing affinity around 4-fold (30.7 mM) (Fig. 2G, circles). All these site-specific variants of both proteins reduce both the affinity and enthalpic contributions to the interaction, thus verifying the involvement of the respective amino acids in the FusB-ferredoxin interface.

### The FusA-FusB interaction requires a salt bridge between FusB R241 and FusA D53

Having elucidated the dimerization and ferredoxin-binding interfaces of FusB at the molecular level we aimed to gain further insight into the FusB-FusA interaction. As mentioned above, the fold of the FusB-CTD monomer in the homodimer structure is most closely related to *E. coli* TonB, based on DALI fold analysis. In the TonB-BtuB co-crystal structure^23^, the TonB-box (DTLVVTAN) of the receptor aligns with residues 225-231 of the terminal b-strand of TonB, with additional interactions mediated by TonB R156, Q158 and Q166. The extreme C-terminal region of TonB (6 residues) appears disordered and is not resolved in the structure. Notably, TonB R156, located next to the highly conserved QPQYP motif, forms a salt bridge with D26 of the receptor. An overlay of FusB-CTD homodimer and TonB bound to BtuB shows a remarkable and surprising similarity: the C terminus of one FusB-CTD protomer appears to partially mimic the TonB-box of BtuB (Fig. 3A), and the intermolecular FusB-CTD R241-D322 salt bridge overlaps directly with the salt bridge between TonB and BtuB. As (i) the predicted FusB-box (DTILVRST) of FusA is very similar to the TonB-box of BtuB, with FusA D53 in an equivalent position to BtuB D26, and (ii) FusB-CTD shows high sequence and fold similarity to TonB-CTD, it seemed likely that residues 307-313 of FusB (analogous to TonB 225-231) would interact with the FusB-box of FusA. We therefore hypothesised that the FusB-box of FusA binds to one molecule of FusB, displacing the long C-terminal β-strand of the other FusB molecule (homodimer state) or the fourth β-strand of the same protomer (monomer state). This is concomitant with the disruption of the inter- or intramolecular salt bridge, whereby R241 would switch from D322 of FusB to D53 of FusA, and the FusB-box of the receptor would align against residues 307-313 located within the long terminal β-strand of FusB. The extreme C-terminal tail of FusB possibly remains unstructured, as in the case of BtuB-bound TonB.

**Figure 3.**
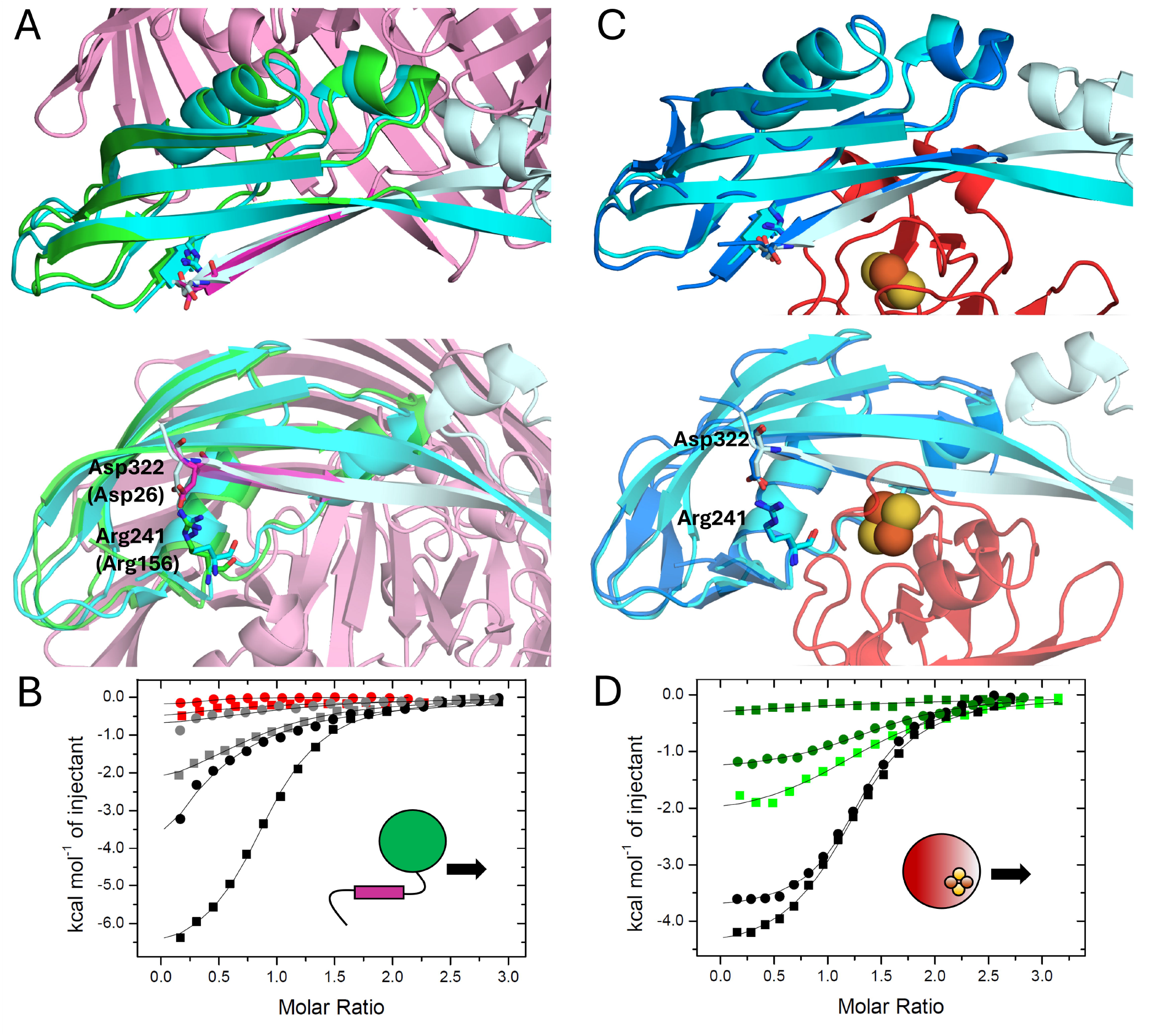
FusB R241 acts as a molecular switch that defines different stages of ferredoxin import. A – overlay of FusB-CTD homodimer (cyan and pale cyan) onto *Ec*TonB-CTD (green) in complex with BtuB (magenta). The TonB-box of the receptor is shown in dark magenta, and the residues involved in salt bridge formation (TonB R156, BtuB D26, FusB R241 and D322) are shown as sticks. The bottom view is a 90° rotation around the horizontal axis. B - ITC thermograms from titrations of FusA_NTR_-GFP wild type (black symbols) or D53A (grey symbols) into FusB-CTD wild type (circles) or D322N (squares) as well as FusA-ferredoxin competition assays (red symbols), whereby FusA_NTR_-GFP was injected into the cell containing FusB-CTD with 2.5-molar excess of SpFer. C – overlay of FusB-CTD homodimer onto monomeric FusB-CTD (dark blue) in complex with ferredoxin (red). R241 and D322 are shown as sticks and the iron-sulfur cluster as spheres. D - ITC thermograms from titrations of SpFer wild type into FusB-CTD only (black symbols, circles – wild type, squares – D322N), or in the presence of 2.5 molar excess of FusA_NTR_-GFP wild type (dark green). The competition assay of FusB-CTD D322N was also performed with 2.5 – molar excess of FusA_NTR_-GFP D53A (light green).

We have previously demonstrated that FusB-CTD interacts with the GFP fusion construct containing the N-terminal region (residues 21-66) of FusA (FusA_NTR_-GFP) with mid-micromolar affinity^11^ (K_D_ = 31.2 µM). To verify whether FusB R241 forms an ionic interaction with FusA D53 we performed a series of ITC experiments, using wild-type constructs and the three aforementioned FusB-CTD variants (R241K, R242K and D322N). In addition, we generated the construct FusA_NTR_-GFP D53A as well as a non-polar R241A variant of FusB-CTD, which, like R241K and D322N, was entirely monomeric (Fig. S2). Curiously, titration of wild-type FusA_NTR_-GFP into wild-type FusB-CTD or the R242K variant (both dimerisable) always produced a hyperbolic-shaped thermogram that was often impossible to fit with 1:1 binding stoichiometry, whereas titrations into the monomeric variants resulted in more typical thermogram shapes (Fig. S3). In fact, binding to D322N variant produced a sigmoidal curve with a low-micromolar affinity (K_D_ = 9.4 µM) and a much larger enthalpic contribution than any interaction we have observed (Fig. S3 and 3B) – including FusB-SpFer binding. These data suggest that that the heats resulting from FusA_NTR_-GFP titration into dimerisable FusB-CTD constructs represent both dissociation of the carboxyl termini of FusB (bound in *trans* or in *cis*), concomitant with disruption of the R241-D322 salt bridge, as well as the competitive binding of FusA_NTR_-GFP. Therefore, FusB-CTD D322N is an entirely monomeric version of otherwise “wild-type” behaving FusB-CTD, with the crucial R241 remaining free and fully available to form a salt bridge with FusA D53. The absence of D322 effectively enhances FusB complexation with FusA_NTR_-GFP (Fig. S4).

Importantly, the interaction of the two R241 variants with FusA_NTR_-GFP produced very low heat, which strongly suggests that this residue is involved in binding FusA. The seemingly conservative lysine substitution provides only a marginal increase in affinity and binding enthalpy (Fig. S3, triangles, K_D_ = 25.3 mM) compared to the alanine variant (inverted triangles, K_D_ = 38.8 mM). Finally, we titrated FusA_NTR_-GFP D53A into FusB-CTD wild-type (Fig. 3B, grey circles, K_D_ = 31.6 mM) and D322A (grey squares, K_D_ = 31.8 mM), which in both cases produced significantly lower heat compared to wild-type FusA_NTR_-GFP titrations (black symbols). Interestingly, however, despite the drastic decrease in the enthalpic contribution, the apparent affinity of FusA_NTR_-GFP D53A for FusB-CTD D322N, or FusA_NTR_-GFP wild type for R241 variants of FusB, was reduced only ∼3-fold, indicating that the FusB-box of FusA still interacts with FusB via strand addition even in the absence of the FusA-FusB salt bridge. The above data support the model whereby FusA_NTR_ aligns against the initial stretch of the long C-terminal β-strand of FusB, displacing the carboxyl terminus of the same FusB molecule or its homodimer partner. This displacement is concomitant with FusB R241 salt bridge switching from FusB D322 (in *trans* or in *cis*) to FusA D53.

### Ferredoxin outcompetes FusA bound to FusB and promotes the formation of R241-D322 salt bridge in *cis*

Once FusB-mediated remodelling of FusA plug is complete, and ferredoxin appears within the receptor lumen, FusB interacts with the substrate to finalise its import. This suggests that ferredoxin needs to outcompete the FusB-box of FusA that is bound to FusB. The structure of the FusB-ferredoxin complex shows that the substrate interacts primarily with two regions of FusB-CTD: the unstructured N-terminus (containing R241 and R242), constituting an interaction hotspot, and the C-terminal β-strand (residues 314-324). The fourth β-strand, which forms the majority of ferredoxin-binding interface, results from the extended C-terminal strand in the FusB homodimer structure forming a hairpin, stabilised by an intramolecular R241-D322 salt bridge. An overlay of FusB-ferredoxin complex onto the FusB homodimer (Fig. 3B) shows that the fourth β-strand aligns against FusB residues 307-313, implying it is in direct competition with the carboxyl termini of the other FusB protomer (see the homodimer structure) or the FusB-box of the receptor (see FusB-TonB overlay, Fig. 3A). The initial stretch of the third β-strand therefore serves as the second interaction hotspot within FusB.

Ferredoxin seems to readily bind FusB regardless of its oligomeric state and the presence of R241-D322 salt bridge, judging from indistinguishable substrate binding parameters for titrations into FusB-CTD wild type and D322N. FusA_NTR_-GFP also interacts with FusB, although less strongly in the context of the wild type protein whereby FusA D53 needs to compete against D322 for access to R241. We also decided to probe the potential competition for FusB between FusA and ferredoxin. To this end we performed an ITC assay with sequential titrations of FusA_NTR_-GFP and SpFer, in either order, into FusB-CTD wild type or D322N variant. This means that in the second titration in the assay protein X is injected into the cell containing FusB-CTD in the presence of ∼2.5 molar excess of protein Y. Titration of SpFer into FusA_NTR_-GFP produced only heats of dilution indicating that these constructs do not interact with each other (Fig. S5). When FusA_NTR_-GFP was injected into the cell containing either construct of FusB-CTD pre-titrated with SpFer, no heats of binding were detected, suggesting that, in the presence of ferredoxin, FusA_NTR_-GFP cannot interact with FusB-CTD (Fig. 3B, red symbols). Injection of SpFer into the cell containing FusB-CTD wild-type pre-titrated with FusA_NTR_-GFP yielded significant heats, implying that SpFer can displace FusA bound to FusB-CTD, yet the apparent affinity and binding enthalpy were reduced (Fig. 3D, dark green circles) compared to the titration into FusB-CTD alone (black circles). Affinity change appeared to be approximately 3-fold lower (K_D_ = 22.3 µM), although this may not be a realistic estimate given that different phenomena (binding/dissociation) are taking place in the reaction cell.

Curiously, when the assay was performed on FusB D322N, SpFer injection produced no discernible signal (Fig. 3D, dark green squares), implying that ferredoxin cannot compete against FusA_NTR_-GFP. However, SpFer could displace FusA_NTR_-GFP D53A variant (light green squares), which, as we demonstrated before, interacts with FusB-CTD D322N with lower affinity as it cannot form the crucial salt bridge with FusB R241. Given that FusB-CTD D322N is also unable to form a salt bridge with R241, both in *trans* and in *cis*, it can outcompete FusA D53A, however it cannot disrupt the already formed salt bridge between FusA D53 and FusB R241.

In summary, ferredoxin and FusA compete for the same region of FusB, with ferredoxin capable of displacing the N-terminal portion of the FusA receptor. The FusB D322N mutation does not affect ferredoxin binding while it enhances the interaction of FusB with FusA by providing full access to FusB R241. However, once FusA is bound to FusB-CTD D322N, the substrate can no longer outcompete FusA: the FusA-FusB salt bridge cannot be broken without FusB D322 displacing FusA D53. This precludes the folding of the C-terminal β-strand of FusB and thus the formation of the substrate-binding interface. These results support the predicted sequence of events, whereby after removing the receptor plug, FusB switches towards the ferredoxin to facilitate its final import step, and FusB R241 acts as a molecular switch.

### The FusB-CTD sequence is highly conserved within *Pectobacterium* genus

We investigated the conservation level of FusB-CTD to establish whether our findings, especially the molecular switch mechanism, are applicable to all Fus systems. Although putative Fus homologues have been found in distantly related bacterial species^28^ they may not be linked to ferredoxin import and we therefore focussed on *Pectobacteriaceae*. Within this family Fus systems are predominantly found in two genera, *Pectobacterium* and *Dickeya*, along with a *Brenneria goodwinii* as well as *Samsonia erythrinea* – a plant pathogen assigned to *Yersiniaceae* family, yet recently proposed to belong to *Pectobacteriaceae*^29^. Based on the alignment of consensus sequences against *Pc*LMG2410 FusB (Fig. 4A), the double arginine motif is completely conserved across all the sampled species, whereas the C-terminal aspartate appears to be conserved solely in *Pectobacterium* and *Samsonia* – it is absent from virtually all other FusB sequences. In fact, the C-terminal end of FusB appears to be generally quite divergent across the different genera. This includes one substrate-binding residue, *Pc*LMG2410 FusB R319, which is substituted with a glutamine (*Brenneria*/*Samsonia*) or glutamate (*Dickeya*) - the latter being a complete charge inversion. Curiously, comparison of the FusB conservation and consensus between *Pectobacterium* and *Dickeya* (Fig. 4B) indicates that pectobacterial FusB sequences are highly similar whereas *Dickeya* species exhibit substantial sequence diversification.

**Figure 4.**
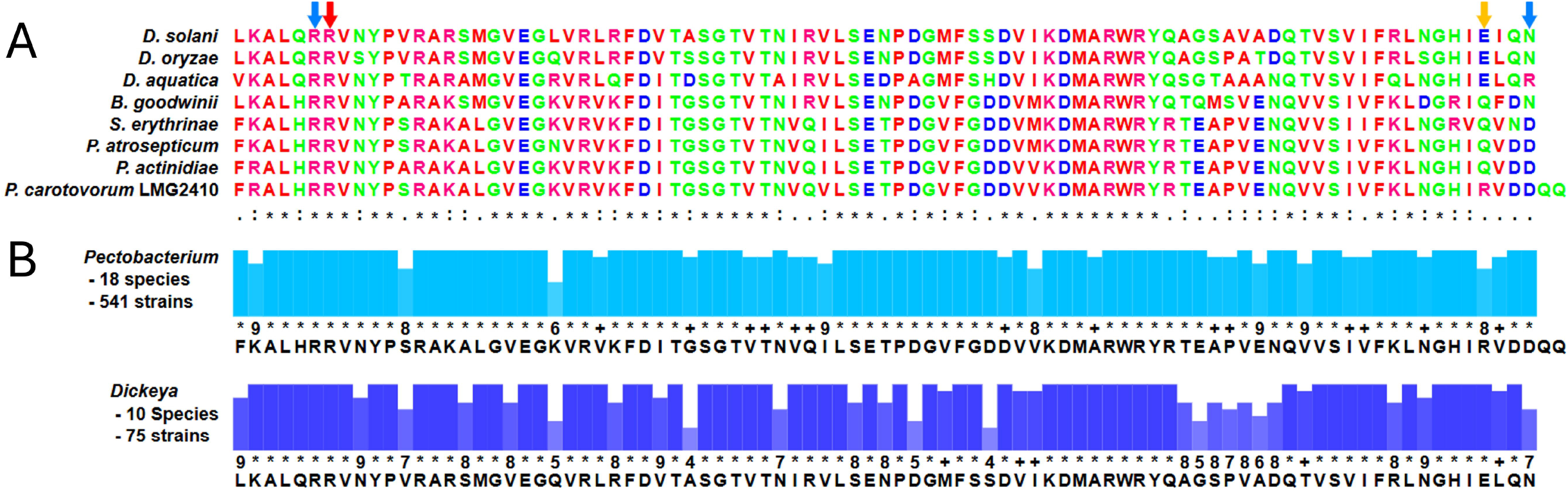
FusB R241-D322 switch mechanism is conserved within *Pectobacterium* genus. A – Multiple sequence alignment of FusB-CTD from *Pc*LMG2410 and related organisms. Conservation status is denoted below each residue: asterisk - fully conserved, colon – conservative substitution, dot – semi-conservative substitution. Blue arrows point at the salt bridge residues, red arrow at R242 (critical for substrate interaction) and orange at R319 which is involved in substrate binding, however the positive charge is lost (*Brenneria*) or reversed (*Dickeya*). B – Conservation and consensus sequence of FusB-CTD within *Pectobacterium* and *Dickeya* genera. Residue conservation is represented by coloured bars with a number or symbol below denoting the level: asterisk – completely conserved, + - conservative substitution, number – range from 1 (lowest) to 9 (highest).

## Discussion

We have previously reported that the Fus-specific TonB-like protein FusB has evolved two distinct roles in the context of importing an unusually large substrate, which required the evolution of a ferredoxin binding interface as well as a process involving two sequential rounds of FusB-mediated force transduction. Plant ferredoxins (∼10 kDa) are comparable in size to the receptor plug (∼15 kDa), which implies that the entire plug needs to be displaced to allow substrate passage. We hypothesise that the substrate interacts with the extracellular face of the plug which would allow it to translocate along the β-barrel concomitantly with FusB-mediated plug remodelling. However, according to results from force measurements on the TBDR BtuB, the plug gradually unravels or separates into subdomains of varying force lability^14^, which suggests that the plug-ferredoxin interaction may be disrupted at some point. Once the plug is dislodged the substrate may therefore need an extra final pull into the periplasm, which would explain its interaction with FusB. Even though the globular domain of FusB largely resembles a canonical TonB-CTD (including the “generic” TonB of *Pc*LMG2410^11^), the polyproline linker appears longer (by 36 – 44 residues), which is consistent with the model whereby FusB contacts the ferredoxin within the receptor lumen.

Dimerisation is a common feature for CTD constructs of most TonB proteins, yet its biological significance has been a controversial matter and only monomeric TonB can bind to the receptor ^23,30–32^. FusB-CTD exists in a monomer-dimer equilibrium in solution and our data indicate that both states (and the respective *cis* and *trans* R241-D322 salt bridges) can be disrupted by the FusB-box of FusA emerging in the periplasm. We propose that under resting conditions FusB is a dimer, given the conformational similarity between FusB protomers in the homodimer structure and the predicted receptor-bound FusB. However, the cellular FusB concentration may not reach the dimerisation-inducing concentrations used *in vitro* and FusB-CTD homodimer may well be an artefact - although we cannot rule out that other parts of FusB protein contribute to dimer formation *in vivo*. Furthermore, FusB molecules may co-localise in the membrane, mirroring the clustering of outer membrane proteins, which have previously been shown to form organised assemblies^33,34^. Co-localisation would locally increase FusB concentration, facilitating homodimerization. Regardless of the oligomeric state it is likely that two FusB molecules would be necessary during the import process as one needs to hold the receptor plug away from the lumen to allow the substrate to emerge on the periplasmic side. According to a recently postulated model TonB-like proteins may move laterally within the inner membrane while transducing force, far away from the uptake site^16^, which further supports the engagement of two FusB molecules. Importantly, as soon as the substrate appears, any FusB near the import site needs to preferentially bind to ferredoxin to complete the import process.

Taking all our results into account we propose a model for Fus-mediated ferredoxin import, presented in Figure 5. In the resting state, FusB-CTD exists as a homodimer (1), with the R241-D322 salt bridge *in trans* stabilising the interaction between one monomer and the long C-terminal β-strand of the other. When ferredoxin binds to the extracellular face of the receptor (2), FusA_NTR_ is released into the periplasm and it displaces the C-terminal β-strand of one FusB molecule within the homodimer, aligning against residues 307-313 of the β-strand of the other FusB (3). Receptor binding involves the R241 salt bridge switch whereby FusA D53 displaces FusB D322. The extreme carboxyl terminus (residues 314-324) of the now monomeric FusB probably remains free and unstructured. FusA plug is then pulled out by FusB with the PMF energy harnessed by ExbBD (4), and the substrate shuffles along the receptor lumen towards the periplasm. Once the ferredoxin appears in the FusA lumen, the long C-terminal β-strand of FusB folds over, forming two shorter strands (5). This conformational change is likely induced by ferredoxin binding – we hypothesise that this brings the extreme C terminus of FusB, including D322, close to the double arginine region. R241 then switches its partner once again, from FusA D53 to FusB D322 (*in cis*), which disrupts FusB-receptor interaction as the newly formed fourth β-strand displaces the FusB-box of FusA. The intramolecular R241-D322 bond stabilises the hairpin which forms the primary substrate-binding interface. FusB guides the substrate into the periplasm by pulling it in the second PMF-energised step (6). Finally, ferredoxin is digested by the protease FusC (7) and the free iron-sulphur cluster is then taken into the cytoplasm via FusD (not shown).

**Figure 5.**
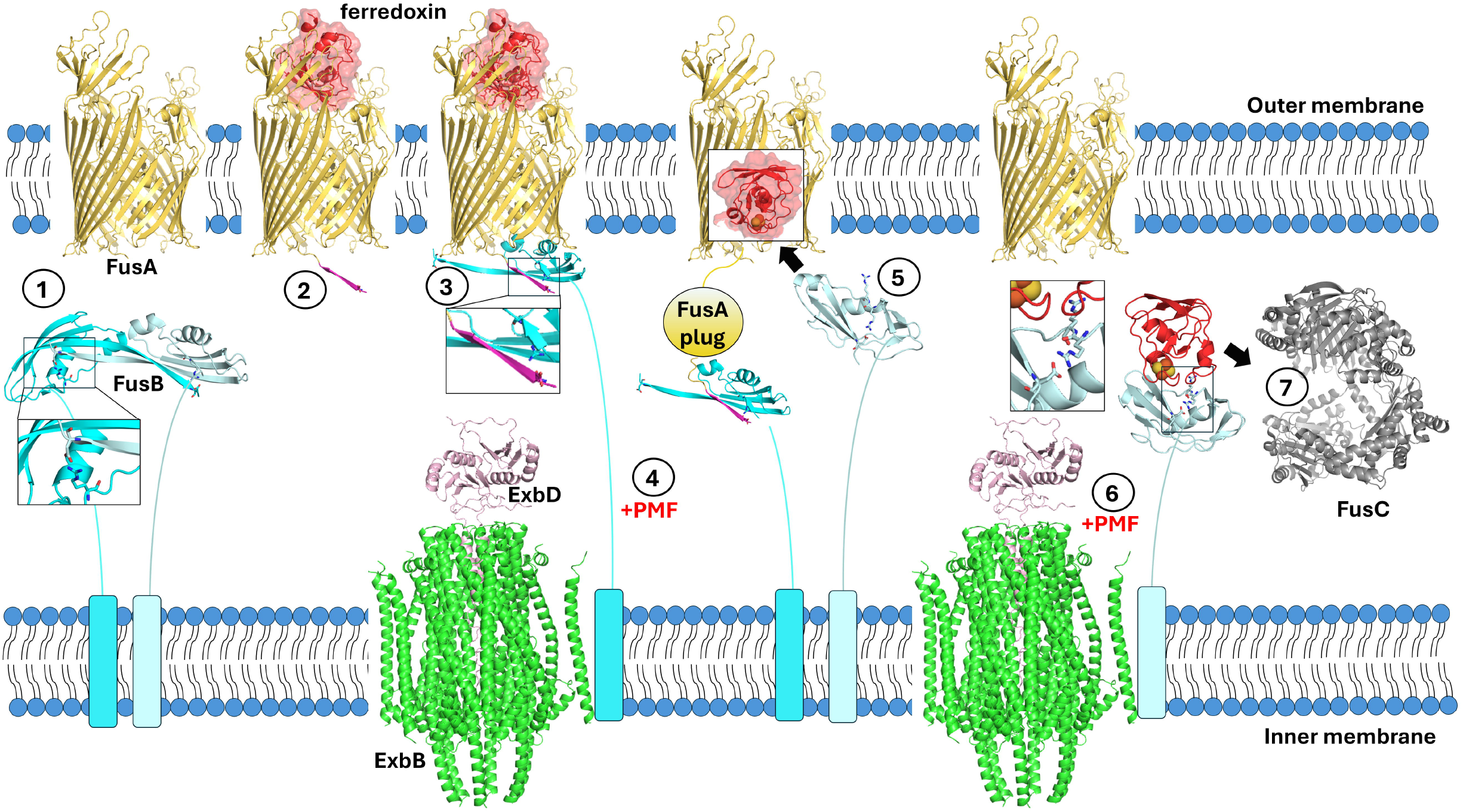
Proposed mechanism for ferredoxin import. A schematic representation of substrate import by Fus, see text for details on the mechanism. All individual structures visualised here are experimental except for (i) FusA-FusB interaction, which was modelled by overlaying onto BtuB-TonB structures, with the FusB-box of FusA represented by TonB-box of BtuB (β-strand in magenta) and the unstructured N-terminal region of the receptor omitted for clarity, and (ii) FusA-ferredoxin complex which was previously generated by docking^21^. The N-terminal extension present in pectobacterial FusBs is not shown. FusB R241, R242 and D322 as well as “FusB-box” aspartate are shown as sticks; FusB R242 is only shown in the context of substrate binding (pale cyan FusB monomer).

We have noticed that the equivalent D322 residue is absent in FusB from other genera, suggesting that our import model may only apply to Fus in *Pectobacterium* spp. and *Samsonia erythrinea*. It could however be reasoned that in some cases ferredoxin(s) may be able to displace the receptor bound to FusB regardless of the receptor-FusB salt bridge, or they simply do not need the extra help from FusB. This could be due to stronger substrate interactions with FusA plug. Furthermore, the receptor-FusB salt bridge may not always be necessary – interaction via strand addition could be sufficient for plug remodelling. Although in the crystal structure TonB forms an ionic bond with BtuB^23^, there does not seem to be any salt bridge between TonB and FhuA^30^, and mutational analysis in *E. coli* demonstrated that none of the arginines at the amino terminus of TonB-CTD are essential for receptor interaction^35^. Further studies will be necessary to verify whether FusB-substrate interaction and R241 molecular switch are essential for all Fus import machineries.

We have hereby gained further insight into the molecular mechanism behind a unique uptake system, which translocates a folded protein across the outer membrane. The process is likely to be energetically costly, yet the growth enhancement conferred by ferredoxin is substantial^11,20,21^. This suggests that ferredoxins may constitute one of the primary sources of iron (and possibly sulphur and amino acids) during soft rot infection. Elucidation of ferredoxin uptake mechanism may help with understanding the general process of TonB-dependent nutrient acquisition, which constitutes a prominent aspect of bacterial virulence. Given that Fus is parasitised by natural protein antibiotics, it may be exploited to devise methods for protecting crops against soft rot pathogens.

## METHODS

### Bacterial strains, media and growth conditions

*E. coli* DH5α cells were used for cloning and BL21 (DE3) for protein production. The cells were incubated at 37°C on LB plates and in LB broth, except for overexpression of ferredoxins or FusB-ferredoxin fusion whereby 2xYT broth was used. For protein production starter LB broth cultures (grown overnight) were diluted 1/75 in 1L media and the cultures were grown with shaking until optical density at 600nm reached 0.5 – 0.8. LB cultures were then induced with 0.7 mM IPTG and grown for 4 -5 hours at 28 °C, whereas 2xYT cultures were supplemented with 0.3 mM IPTG and grown overnight at 20 °C. All media were supplemented with 100 µg ml^-1^ ampicillin, except those used for cells harbouring pWaldo plasmid (for expression of FusA_NTR_-GFP) which were supplemented with 50 µg ml^-1^ kanamycin.

### Generation of the expression plasmids

Plasmids and primers used in this study are listed in Tables S1 and S2, respectively. Codon-optimised gene strings were used as templates for PCR using designated primers to generate pJ404 ferredoxin expression vectors, with the resulting proteins containing 6His-tags at the C terminus. These plasmids were generated as described before^11^: both pJ404 vector and the PCR amplicon were digested with NdeI and XhoI and then ligated together. All other plasmids were made either by site-directed mutagenesis or by sub-cloning genes (or their portions) and inserting them into desired locations on target vectors using NEBuilder HiFi DNA Assembly kit (NEB). To allow the assembly target plasmids were first linearised by digestion with XhoI (pExp) or XhoI and NdeI (pJ404).

**Table S1.** List of plasmids used in this study.

| PLASMID | SOURCE | PARENT PLASMID |
| --- | --- | --- |
| pJ404FusBCTD | Wojnowska *et al* 2020<sup>11</sup> | pJ404 |
| pJ404FusBCTD-R241K | This work | pJ404FusBCTD |
| pJ404FusBCTD-R241A | This work | pJ404FusBCTD |
| pJ404FusBCTD-R242K | This work | pJ404FusBCTD |
| pJ404FusBCTD-H317A | This work | pJ404FusBCTD |
| pJ404FusBCTD-D322N | This work | pJ404FusBCTD |
| pExpFusBCTD-SpFer | This work | pExp |
| pJ404SpFer | This work | pJ404 |
| pExpNoTevSpFer | This work | pExp |
| pExpTEVSpFer | This work | pExp |
| pWaldoFusANTR-GFP | Wojnowska *et al* 2020<sup>11</sup> | pWaldo |
| pWaldoFusANTR-GFP-D53A | This work | pWaldoFusANTR-GFP |

**Table S2.**
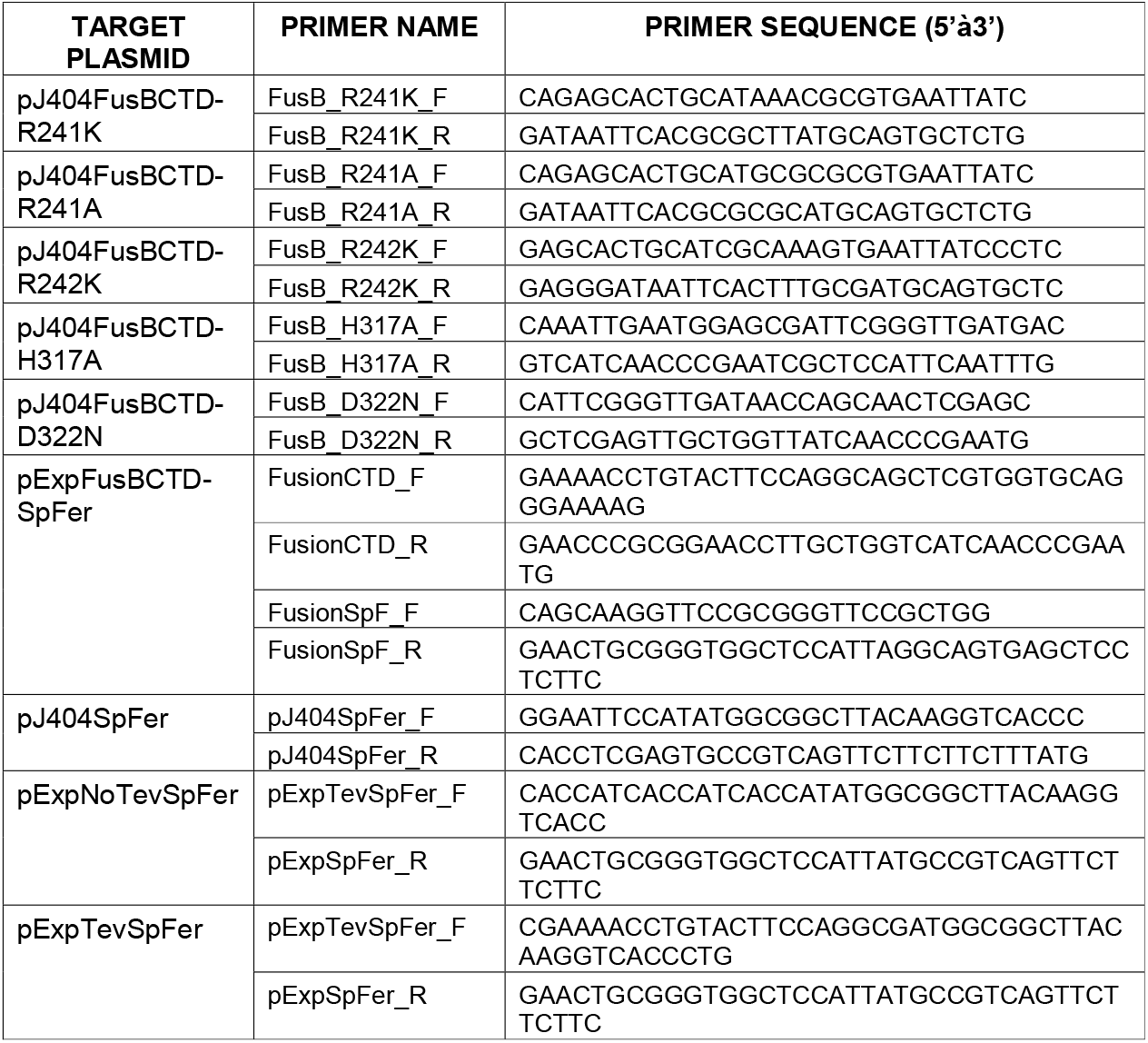
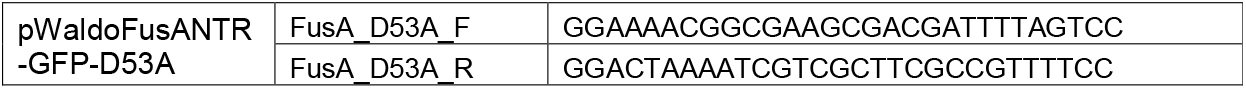
List of primers used in this study.

### Protein purification

FusA_NTR_-GFP, FusC, FusB-CTD constructs and ferredoxin constructs with uncleavable 6His-tags were purified as described before^11,21^. Briefly, pelleted cells were resuspended in Buffer A (50 mM Tris-HCl, 500 mM NaCl, pH 8) supplemented with DNAse (Merck) and a protease inhibitor cocktail tablet (cOmplete, Roche) and lysed by passing through a cell disrupter. After pelleting the debris, the supernatant was loaded onto pre-equilibrated NiNTA resin. The bound protein was washed with Buffer A containing 20 mM imidazole and eluted in Buffer B containing 300 mM imidazole. Eluate was concentrated and loaded onto Superdex 75 Increase gel filtration column (Cytiva) pre-equilibrated in SEC Buffer (20 mM HEPES, 140 mM NaCl, pH 8). SpFer and FusB-ferredoxin fusion with the cleavable N-terminal tags were eluted from NiNTA resin, supplemented with 5 mM DTT (Merck) and TEV protease (produced in-house) at a 1/50 weight ratio and dialysed in a 3 kDa MWCO membrane against 20 mM Tris-HCl, 100 mM NaCl, pH 8 buffer overnight at 4 °C. Digested protein was filtered and passed through NiNTA resin, the flow-through was concentrated and subjected to gel filtration as described above. Fractions containing pure protein, as judged SDS-PAGE, were pooled, concentrated and stored at -80 °C.

### FusC cleavage assays

Digestion reactions were performed at room temperature in 20 mM HEPES, 50 mM NaCl, pH 8 buffer with 30 µM ferredoxin and 4 µM FusC. 15 µl aliquots were taken at specified time points and a control ferredoxin sample without protease was incubated alongside for 150 min. The samples were quenched with 5 µl of 4x loading dye containing 5 mM DTT, boiled for 5 min and the proteins were resolving on a 10% Bis-Tris gel (mPAGE).

### Analytical gel filtration and SEC-MALS

Co-migration assays were conducted on a Superdex 75 Increase column (Cytiva) equilibrated in the SEC Buffer, with chromatograms recorded at 280 and 330 nm. 300 µl sample containing 120 µM FusB-CTD and/or 150 µM ferredoxin was injected per run. SEC-MALS experiments were performed in the same buffer and using Superdex 75 Increase column on an AKTÅ chromatography system (Cytiva) coupled to a multi-angle light scattering detector, DAWN HELEOS II (Wyatt Technology) and Optilab T-rEX refractometer (Wyatt Technology). Data was analysed using Astra 7.3 software (Wyatt Technology).

### Isothermal titration calorimetry

All experiments were performed at 25 °C in SEC Buffer on MicroCal iTC200 instrument (Malvern Panalytical) with differential power set to 3. Typically 1.1 mM analyte was injected 19 x 2 µl into the cell containing 85 µM ligand and titration was continued until only heats of dilutions were observed.

We have previously shown that natively extracted and commercially acquired *Spinacia oleracea* (spinach) ferredoxin 1 (SpFer) binds to FusB-CTD with an affinity in the low micromolar range (K_D_ = 7.8 µM) ^11^, however the recombinant SpFer construct with a C-terminal 6His-tag appeared to bind with two-fold lower affinity and slightly reduced enthalpy (Fig. S6). To test whether the tag interferes with the FusB interaction we generated N-terminally modified SpFer constructs with a 6His-tag or a cleavable 8His-tag. Both N-terminally tagged and native-like tag-free construct exhibited comparable binding parameters to the natively extracted ferredoxin, and were used interchangeably throughout.

In the competition assays one ligand (FusA_NTR_-GFP or SpFer) was titrated into the cell containing FusB-CTD after which the excess of liquid (∼40 µl) was removed and the other ligand was titrated into the cell, which now contained a slightly lowered concentration of FusB-CTD in the presence of ∼2.5 molar excess of the first ligand. Table S3 lists all the titrations, number of repeats and calculated mean thermodynamic parameters determined with MicroCal Origin software.

**Table S3.**
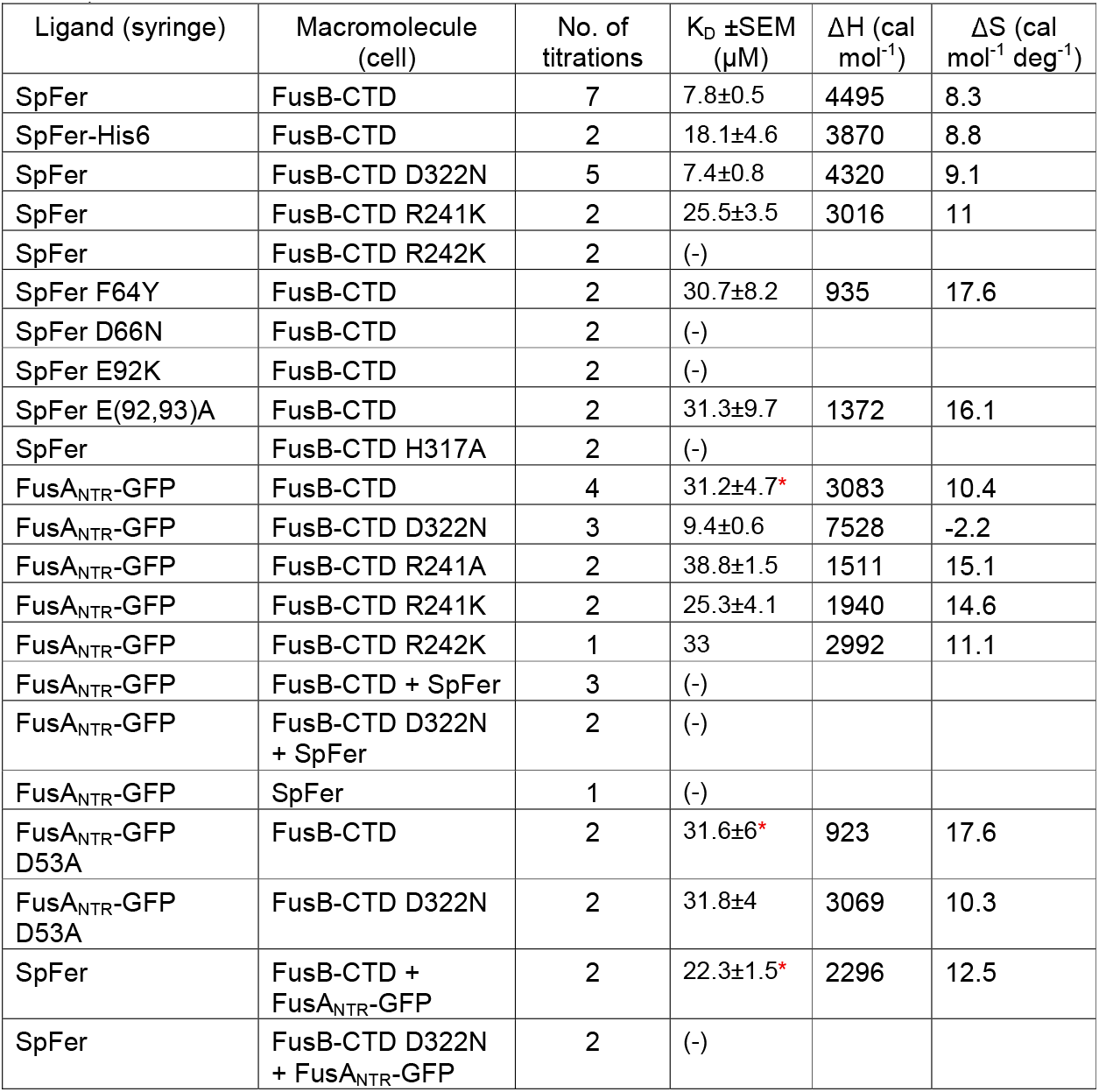
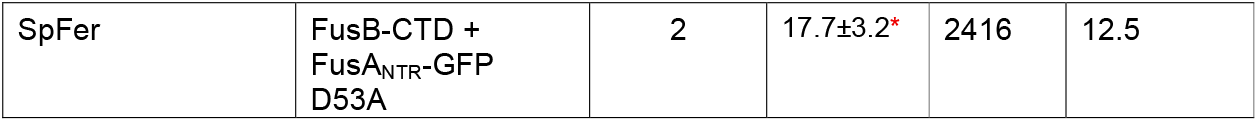
List of all ITC experiment types and calculated thermodynamic parameters. Enthalpy contributions are all negative values. Asterisk denotes titrations that produced difficult to fit/interpret thermograms (very low signal, hyperbolic thermogram or simultaneous association/dissociation events).

| Ligand (syringe) | Macromolecule (cell) | No. of titrations | K<sub>D</sub> ±SEM (µM) | ΔH (cal mol<sup>-1</sup>) | ΔS (cal mol<sup>-1</sup> deg<sup>-1</sup>) |
| --- | --- | --- | --- | --- | --- |
| SpFer | FusB-CTD | 7 | 7.8±0.5 | 4495 | 8.3 |
| SpFer-His6 | FusB-CTD | 2 | 18.1±4.6 | 3870 | 8.8 |
| SpFer | FusB-CTD D322N | 5 | 7.4±0.8 | 4320 | 9.1 |
| SpFer | FusB-CTD R241K | 2 | 25.5±3.5 | 3016 | 11 |
| SpFer | FusB-CTD R242K | 2 | (-) |  |  |
| SpFer F64Y | FusB-CTD | 2 | 30.7±8.2 | 935 | 17.6 |
| SpFer D66N | FusB-CTD | 2 | (-) |  |  |
| SpFer E92K | FusB-CTD | 2 | (-) |  |  |
| SpFer E(92,93)A | FusB-CTD | 2 | 31.3±9.7 | 1372 | 16.1 |
| SpFer | FusB-CTD H317A | 2 | (-) |  |  |
| FusA<sub>NTR</sub>-GFP | FusB-CTD | 4 | 31.2±4.7* | 3083 | 10.4 |
| FusA<sub>NTR</sub>-GFP | FusB-CTD D322N | 3 | 9.4±0.6 | 7528 | -2.2 |
| FusA<sub>NTR</sub>-GFP | FusB-CTD R241A | 2 | 38.8±1.5 | 1511 | 15.1 |
| FusA<sub>NTR</sub>-GFP | FusB-CTD R241K | 2 | 25.3±4.1 | 1940 | 14.6 |
| FusA<sub>NTR</sub>-GFP | FusB-CTD R242K | 1 | 33 | 2992 | 11.1 |
| FusA<sub>NTR</sub>-GFP | FusB-CTD + SpFer | 3 | (-) |  |  |
| FusA<sub>NTR</sub>-GFP | FusB-CTD D322N + SpFer | 2 | (-) |  |  |
| FusA<sub>NTR</sub>-GFP | SpFer | 1 | (-) |  |  |
| FusA<sub>NTR</sub>-GFP D53A | FusB-CTD | 2 | 31.6±6* | 923 | 17.6 |
| FusA<sub>NTR</sub>-GFP D53A | FusB-CTD D322N | 2 | 31.8±4 | 3069 | 10.3 |
| SpFer | FusB-CTD + FusA<sub>NTR</sub>-GFP | 2 | 22.3±1.5* | 2296 | 12.5 |
| SpFer | FusB-CTD D322N + FusA<sub>NTR</sub>-GFP | 2 | (-) |  |  |
| SpFer | FusB-CTD +<br>FusA<sub>NTR</sub>-GFP<br>D53A | 2 | 17.7±3.2* | 2416 | 12.5 |

### Protein crystallisation and structure determination

For all crystallisation attempts the protein was always passed through gel filtration in 10 mM HEPES pH8, 30 mM salt immediately before concentrating and setting up crystal screens. FusB-CTD crystals were obtained from screening using various sparse matrix screens and sitting drop vapour diffusion. The aim was to co-crystallise FusB-CTD with ferredoxin, hence FusB-CTD was mixed with commercially acquired SpFer (Merck) at 1:1.5 ratio and the protein mixture was subjected to gel filtration on Superdex 75 Increase in 10 mM HEPES, 30 mM NaCl, pH 8 buffer. The peak fractions were pooled and concentrated to ∼15 mg ml^-1^, then used in screening with 1:1 and 1:1.5 (protein:liquor) drop composition. Best crystals with a red tint, diffracting to 2.08 Å, were obtained at 4 °C in 100 mM Tris-HCl pH 8.5, 8% PEG8000. Indexing, scaling and merging of the data was performed by the Diamond Light Source (DLS) automatic pipeline xia2 and the structure was solved by PHASER^37^ using *Ec*TonB from various available structures as the search model. Searches using ferredoxin model failed, indicating that the crystals contained only FusB-CTD. The best initial model was obtained using *Ec*TonB from PDB 2GSK^23^ and this was further used for iterative model building in Coot^38^ and refinement using Refmac5^39^. Quality assessment of the model was performed using MolProbity^40^.

Crystals of FusB-ferredoxin fusion were also obtained through sparse matrix screening and the red crystals diffracting to 1.7 Å formed at 20 °C in a Morpheus screen condition composed of 0.05M HEPES, 0.05M MOPS pH 7.5, 2M divalent cations (0.005M Manganese(II) chloride, 0.005M Cobalt(II) chloride, 0.005M Nickel(II) chloride, 0.005M Zinc acetate), 0.1 M monosaccharides (0.2 M xylitol, 0.2 M D-fructose, 0.2 M D-sorbitol, 0.2 M myo-inositol, 0.2 M L-rhamnose monohydrate), 20% PEGMME550 and 10% PEG 20000. The same DLS pipeline and software was used for data processing, model building and quality assessment as described above except that the structure was solved using FusB-CTD and spinach ferredoxin (PDB ID: 1A70) as search models. The statistics for data collection and refinement are shown in Table 1 and structures were deposited with PDB accession codes 7ZC8 and 9HI3. The models of protein structures were generated using PyMOL (Schrödinger, LLC).

**Table 1.**
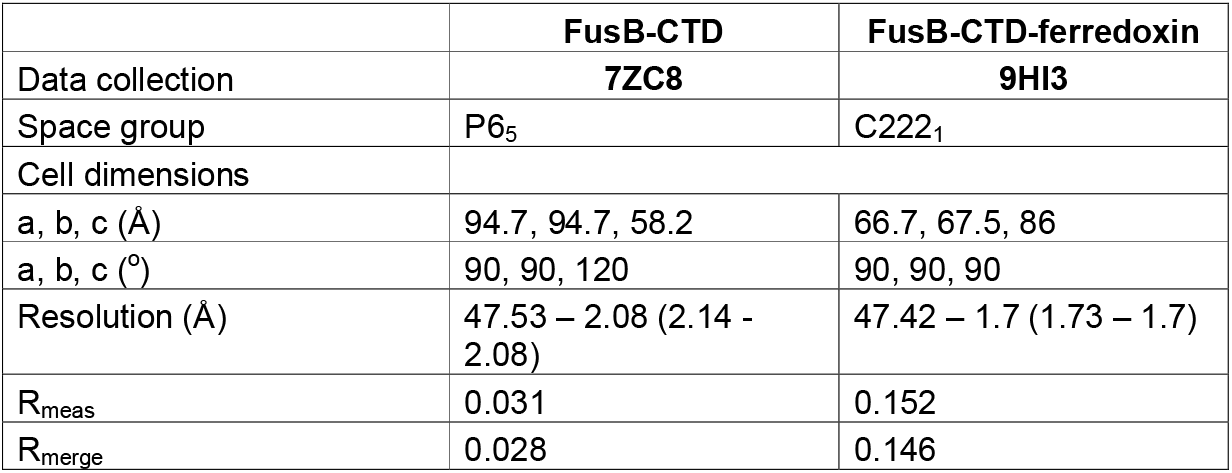
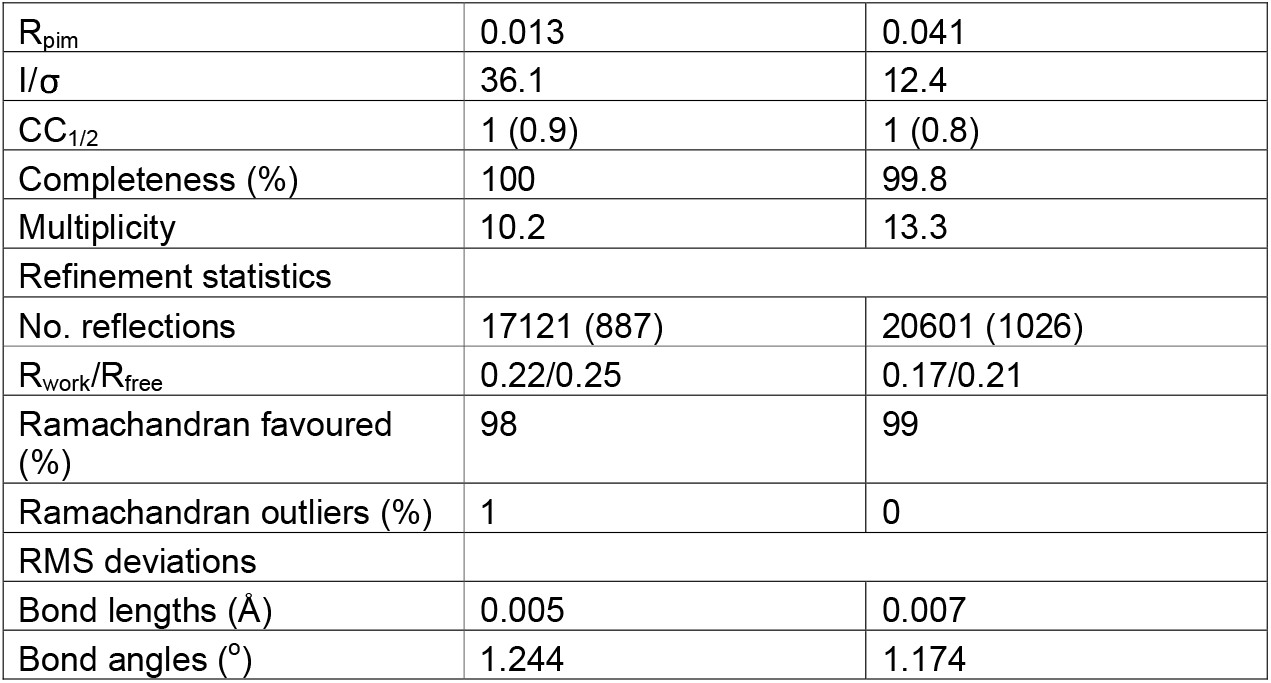
Data collection and refinement statistics.

|  | **FusB-CTD** | **FusB-CTD-ferredoxin** |
| --- | --- | --- |
| Data collection | **7ZC8** | **9HI3** |
| Space group | P6<sub>5</sub> | C222<sub>1</sub> |
| Cell dimensions |  |  |
| a, b, c (Å) | 94.7, 94.7, 58.2 | 66.7, 67.5, 86 |
| a, b, c (°) | 90, 90, 120 | 90, 90, 90 |
| Resolution (Å) | 47.53 – 2.08 (2.14 - 2.08) | 47.42 – 1.7 (1.73 – 1.7) |
| R<sub>meas</sub> | 0.031 | 0.152 |
| R<sub>merge</sub> | 0.028 | 0.146 |
| R<sub>pim</sub> | 0.013 | 0.041 |
| I/σ | 36.1 | 12.4 |
| CC<sub>1/2</sub> | 1 (0.9) | 1 (0.8) |
| Completeness (%) | 100 | 99.8 |
| Multiplicity | 10.2 | 13.3 |
| Refinement statistics |  |  |
| No. reflections | 17121 (887) | 20601 (1026) |
| R<sub>work</sub>/R<sub>free</sub> | 0.22/0.25 | 0.17/0.21 |
| Ramachandran favoured (%) | 98 | 99 |
| Ramachandran outliers (%) | 1 | 0 |
| RMS deviations |  |  |
| Bond lengths (Å) | 0.005 | 0.007 |
| Bond angles (°) | 1.244 | 1.174 |

### Conservation analysis

The genome sequences from 1005 organisms were retrieved from GenBank (Supplementary Table S3), and the amino acid sequences of the encoded proteins were obtained with gffread v0.12.7^41^. FusB and TonB proteins were identified based on the presence of the conserved domains TonB_N (PF16031), TonB_C (PF03544), and TonB_2 (PF13103), which were detected using HMMER v3.4^42^. After removal of sequences of low quality, we kept 619 FusB sequences from *Brenneria* (n=1), *Dickeya* (n=75), *Erwinia* (n=1), *Pectobacterium* (n=541) and *Samsonia* (n=1). To provide a reference, we also analysed 842 TonB sequences belonging to 13 different genera (Supplementary Table S4). The amino acid sequences were aligned using MAFFT v7.526^43^. Construction of consensus sequences and conservation analysis were carried out using Jalview v2.11.4.0^44^.

## COMPETING INTERESTS STATEMENT

The authors declare no competing interests in relation to this work.

## ACKNOWLEDGEMENTS

We thank Professor Ben Luisi for hosting MW and providing valuable guidance and comments, Dr Aleksei Lulla for pEXP and TEV protease expression vectors,

Diamond Light Source and Dr Paul Brear for assistance with crystal data collection and processing. We also thank Broodbank Research Fellowship funders for the Fellowship awarded to MW.

## Notes

### Competing Interest Statement

The authors have declared no competing interest.

