## Supplemental Information for "Receptor-substrate competition for the TonB homologue FusB suggests a model for ferredoxin import in *Pectobacterium* spp"

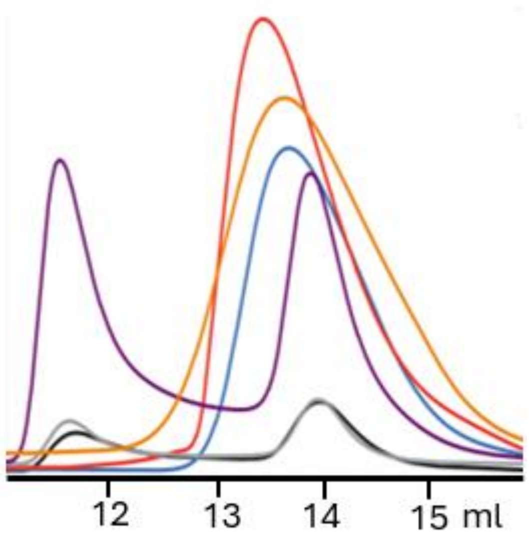


**Figure S1**. Size exclusion chromatograms of FusB-CTD wild type (black line), R242K (grey), H317A (purple), R241K (red), R241A (orange), D322N (blue). 0.5 mg (wild type, R242K) or ~5mg protein (H317A, R241K, R241A, D322N) was loaded.

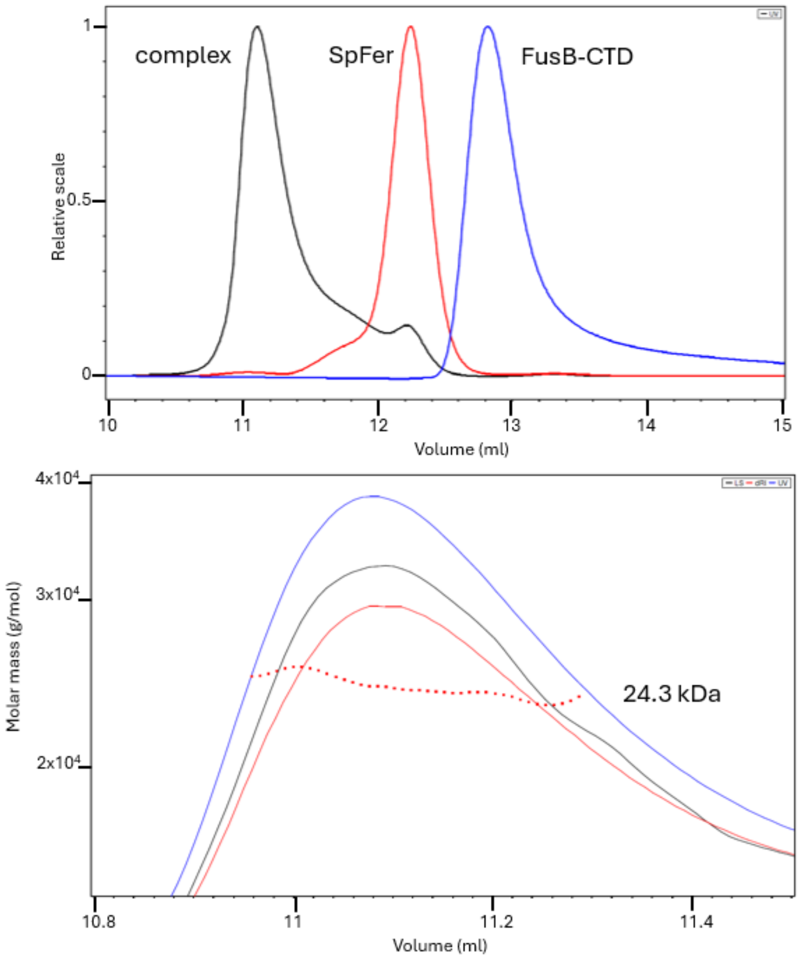


**Figure S2**. SEC-MALS data for FusB-CTD D322N in complex with SpFer. FusB-CTD variant was used to simplify the chromatogram. Top: overlay of UV chromatograms. Bottom: complex peak with molar mass (dotted line). Blue curve – UV absorption, red – refractive index, black – light scattering.

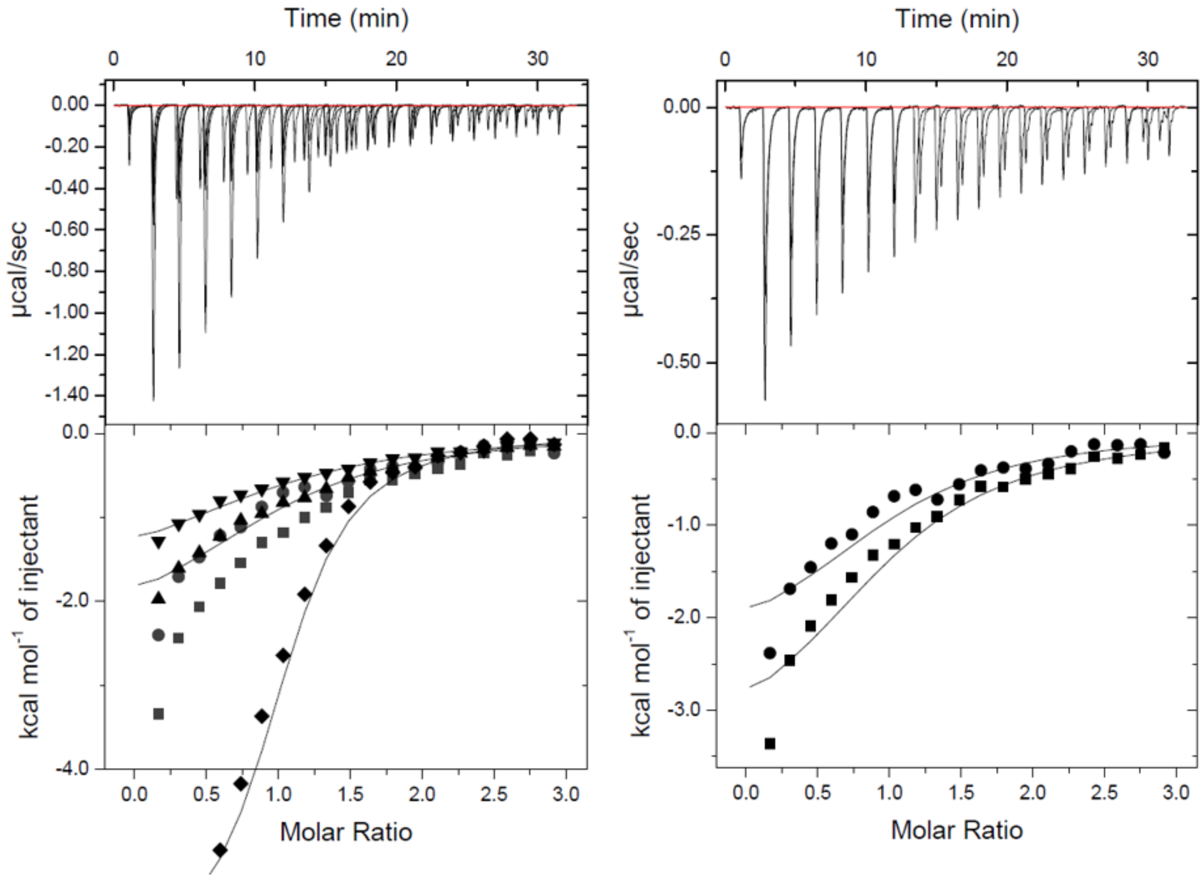


**Figure S3.** Titration of FusA$_{NTR}$-GFP into FusB-CTD wild type (squares), D322N (diamonds), R242K (circles), R241A (inverted triangles) and R241K (triangles). The thermogram for D322N titration is also shown in Figure 4, here it is truncated to better visualise the other thermograms. Note that in the case of wild type and R242K (dark grey on the left panel) the resulting thermograms were hyperbolic in shape, and fitting resulted in stoichiometry parameter N deviating below 0.2. The right panel shows the non-optimal fits for wild type and R242K titrations with N fixed at 1.

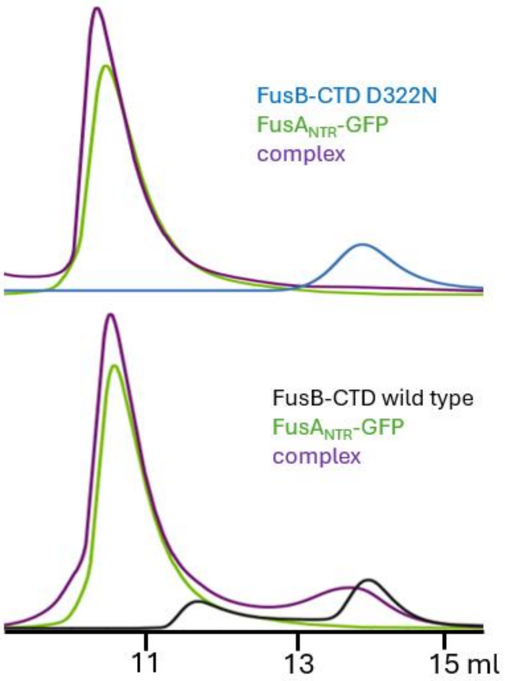


**Figure S4**. Size exclusion chromatograms showing the level of complexation of FusB-CTD wild type (black) and D322N (blue) with FusA$_{NTR}$-GFP (green). In the presence of FusA$_{NTR}$-GFP the peak corresponding to D322N variant disappears completely, whereas wild type peak largely remains, indicating only partial complexation.

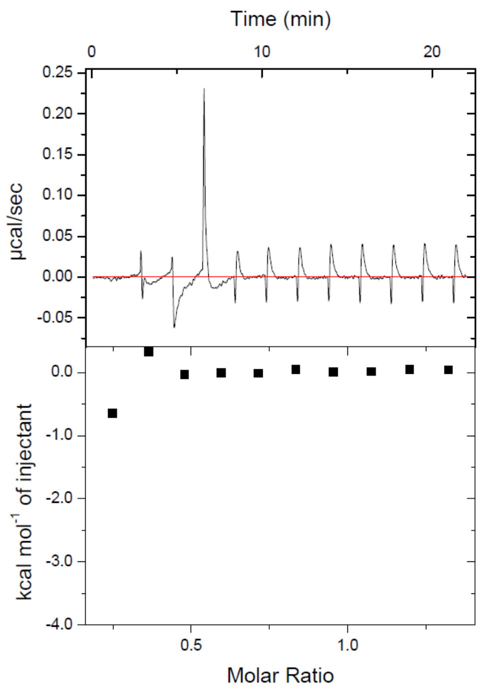


**Figure S5**. Titration of SpFer into $FusA_{NTR}$-GFP.

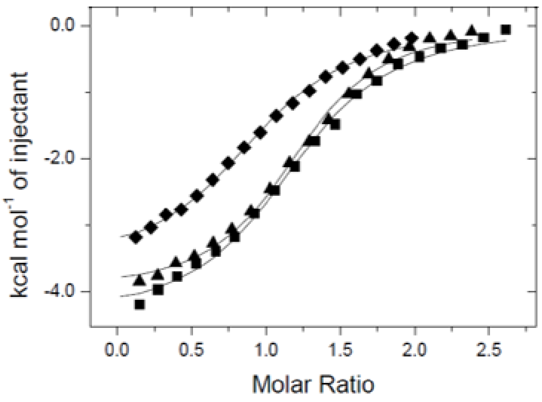


**Figure S6**. Thermograms from the titration of tag-free (squares), N-terminally-tagged (triangles) and C-terminally tagged SpFer (diamonds, $K_D$ = 18.1±4.6 μM) into *Pc*FusB-CTD. Titrations of tag-free and N-terminally tagged SpFer were virtually indistinguishable, with the average $K_D$ = 7.8±0.5 μM.
